# Survey of hippocampal responses to sound in naïve mice reveals widespread activation by broadband noise onsets

**DOI:** 10.1101/2025.04.15.649017

**Authors:** James Bigelow, Toshiaki Suzuki, Yulang Wu, Ying Hu, Andrea R. Hasenstaub

## Abstract

Recent studies suggest some hippocampal (HC) neurons respond to passively presented sounds in naïve subjects, but the specificity and prevalence of these responses remain unclear. We used Neuropixels probes to record unit activity in HC and auditory cortex (ACtx) of awake, untrained mice during presentation of diverse sound stimuli. A subset of HC neurons exhibited reliable, short-latency responses to passive sounds, including tones and broadband noise. HC units showed evidence of tuning for tone frequency but not spectrotemporal features in continuous dynamic moving ripples. Across sound types, HC responses overwhelmingly occurred at stimulus onset; they quickly adapted to continuous sounds and did not respond at sound offset. Among all sounds tested, broadband noise was by far most effective at driving HC activity, with response prevalence scaling with increasing spectral bandwidth and density. Responses to noise were also far more common than visual flash stimuli. Sound-evoked face movements, quantified by total facial motion energy (FME), correlated with population-level HC activity, but many individual units responded regardless of movement, indicating both auditory and motor-related inputs. These results show that abrupt, acoustic events are sufficient to activate HC neurons in the absence of learning or behavioral engagement. This suggests a possible role for HC in detecting salient environmental changes and supports the idea that auditory inputs contribute directly to HC function. Given emerging links between hearing loss and dementia, these findings highlight a potential pathway by which auditory deafferentation could impact cognitive health.

## Introduction

Hippocampus (HC) has a well-established role in processing behaviorally important sounds. For instance, trace conditioning studies demonstrate HC is necessary for bridging the temporal gap between a tone and subsequent shock (McEchron et al., 1998). In tone discrimination learning, HC responses begin to distinguish frequencies predictive of shock or safety in parallel with behavioral discrimination (Freeman et al., 1996). Many other studies have observed sound-related activity in HC during auditory tasks, which is often absent or sharply diminished during passive playback (Aronov et al., 2017; Kumar et al., 2016). These studies suggest the primary role of HC in processing sound may be in learning contingencies and orchestrating behavioral responses.

Nevertheless, several recent findings indicate some HC neurons respond to passively presented sounds even in untrained subjects (Billig et al., 2022; Bimbard et al., 2023; Martorell et al., 2019; Xiao et al., 2018). Such responses are observed at sound levels well below startle and pain thresholds and typically have short latencies (∼15–25 ms). They have been observed using a variety of sounds including broadband noise, tone pips, and natural sounds. One study noted units responsive to passively presented sounds were largely distinct from units that responded to the same sounds during task engagement (Aronov et al., 2017). Thus, auditory inputs are capable of evoking HC responses outside of learning and goal-directed behavioral contexts.

Few studies have attempted to identify specific acoustic features that drive HC responses in passive settings. Sound level is clearly important, considering HC noise thresholds are roughly 30 dB higher than auditory cortex (ACtx). One study found substantially more HC units were activated by broadband noise than pure tones (Xiao et al., 2018), consistent with a similar dissociation in upstream entorhinal cortex, medial septum, and pontine nuclei (G.-W. Zhang et al., 2018). Little else is known about the passive environmental sound features that evoke HC responses. It is also unknown whether and how these responses might complement processing in canonical auditory pathway stations. Thus, their prevalence and functional significance are unclear.

Clarifying the nature and extent of passive auditory processing in HC could provide novel insights into both HC function and hearing, and would be directly relevant to interpreting recent findings suggesting hearing manipulations can modify HC structure and function (Martorell et al., 2019; Qian & Ricci, 2020; Y. Zhang et al., 2021). Among these, multiple studies have found hearing loss induced by noise overexposure or cochlear ablation causes HC dysfunction in rodents, including dendritic simplification, suppressed neurogenesis, and spatial learning and memory deficits (Nadhimi & Llano, 2021; Qian & Ricci, 2020). These findings are significant considering they may be related to the epidemiological link between hearing loss and dementia in humans (Griffiths et al., 2020; Nadhimi & Llano, 2021). The mechanistic cascade linking hearing loss and HC dysfunction is not well understood, but several studies controlling for indirect effects of stress or vestibular pathology support auditory deafferentation as a causal factor. It follows that deafferenting inputs mediating responses to passive environmental sounds could contribute to HC changes to the extent that these inputs drive HC activity.

The current study was motivated by two primary questions. First, which aspects of the auditory environment evoke HC responses in naïve subjects, and how prevalent are these responses? Second, how do responses in HC compare with canonical auditory pathway stations? We addressed these questions in a series of experiments examining a wide range of sound features using Neuropixels probes to simultaneously record from large unit samples in HC and ACtx.

## Materials and Methods

### Subjects and surgery

All procedures were approved by the Institutional Animal Care and Use Committee at the University of California, San Francisco. Subjects were adult male and female C57BL/6 mice (sample sizes given for each experiment below). Most expressed optogenetic effectors targeting interneuron subpopulations, and some were wildtype littermates of 5XFAD transgenics (genotyping performed by Transnetyx). All mice were housed socially under a 12H-12H light-dark cycle. Surgeries were performed under isoflurane anesthesia with perioperative monitoring and analgesics (lidocaine, meloxicam, and buprenorphine). A headbar was implanted above the right temporal lobe with dental cement, after which subjects were allowed to recover at least two days. A small craniotomy (∼1–2 mm diameter) was then made above ACtx (∼2.5–4 mm posterior to bregma) and sealed with silicone elastomer (Kwik-Cast, World Precision Instruments). Electrophysiological recordings were typically conducted within 1–3 days following craniotomy and occasionally the day of the craniotomy following 3+ hours recovery.

### *In vivo* electrophysiology

All experiments were conducted inside a dark sound attenuation chamber (Industrial Acoustics Company). Subjects were headfixed on a spherical treadmill permitting free movement or rest (**Figure 1A**; Bigelow et al., 2022; Niell & Stryker, 2008). Extracellular recordings were made with Neuropixels probes lowered ∼3500 μm below the brain surface (1 μm/s) in a trajectory spanning ACtx, HC, and thalamus (**Figure 1B**, left). Prior to recording, the craniotomy was filled with 2% agarose and allowed to settle for 20+ minutes. Neuropixels data were recorded using OpenEphys or SpikeGLX. Spike sorting was performed offline using KiloSort 2.0 (Pachitariu et al., 2024). We used physiological features to estimate the boundaries of ACtx and HC offline, including depth-dependent transitions in local field potential power, multiunit and single firing rates, unit isolation density, and sound-evoked responses (**Figure 1C**). In some recordings, we applied Di-I to the probe for histological visualization of the probe trajectory (**Figure 1B**, right). Although we often recorded from putative auditory thalamus (units below HC with short latency, tuned responses to tones), we focused analysis on ACtx as recordings more consistently targeted this area and its boundaries were less ambiguous (e.g., unit isolation gaps reflecting pia and white matter). A minority of recordings included only ACtx or HC (e.g., anterior sites hitting ACtx but missing ventral HC). These were included in group data analyses since our primary aim was to define global differences between regions. Spike waveforms for example units are shown throughout as median ±MAD across all recorded spikes with 0.5 ms scale bars.

**Figure 1.**
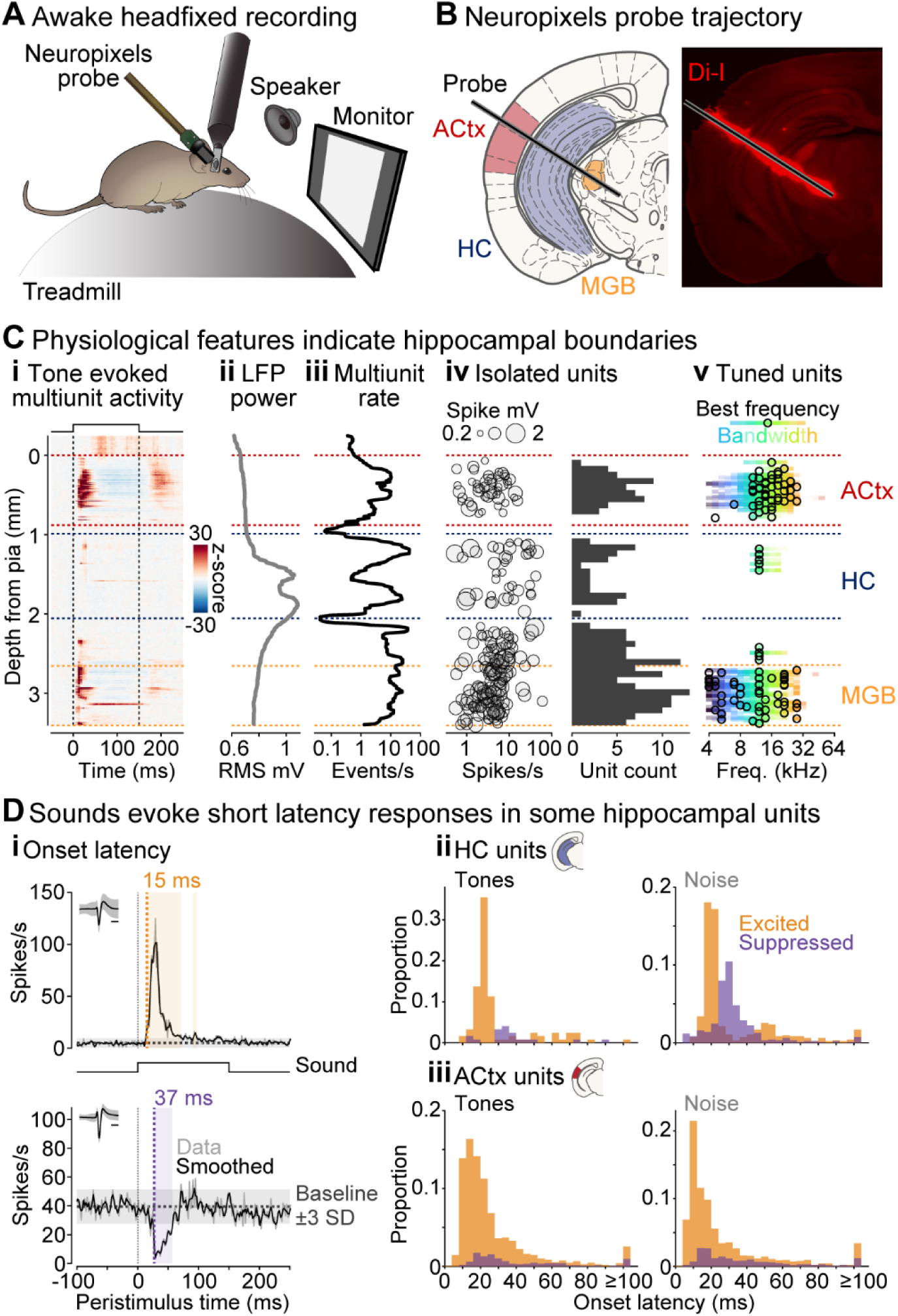
Passive sounds evoke short latency responses in some hippocampal units of naïve mice. (**A**) Sounds were presented passively to naïve mice headfixed on a spherical treadmill. (**B**) Neuropixels probes enabled simultaneous recording in HC and ACtx. (**C**) Multiple physiological features were used for offline estimation of HC and ACtx boundaries including depth dependent transitions in (**i**) tone evoked multiunit activity, (**ii**) LFP power, (**iii**) multiunit event rates, (**iv**) unit isolation density, and (**v**) prevalence of tuned units. (**D**) Units in HC and ACtx. (**i**) Example HC units showing onset latency calculation. (**ii**) Summary of response latencies in HC units for tones (left) and (noise). (**iii**) Summary of response latencies in ACtx units as in (iii).

### Auditory and visual stimuli

All stimuli were generated and presented with MATLAB (Mathworks). Sounds were sampled at 192 kHz and presented with an OctaCapture soundcard (Roland) through an open-field speaker (MF1 or ES1, Tucker-Davis Technologies) positioned ∼20 cm from the left (contralateral) ear. Except where otherwise indicated, sounds were gated with 5 ms cosine^2^ ramps and presented at 65–70 dB SPL. Sound levels were calibrated using either a Brüel & Kjær Model 4939 microphone (Model 2209 meter), or a combination of Avisoft-Bioacoustics CM16/CMPA and Fifine K669B microphones for frequencies above and below ∼5 kHz, respectively.

Sound types tested in experiments below included the following: *Tones*: sinusoid functions of constant frequency (range: 4–64 kHz); *Broadband noise*: white noise filtered with a passband of either 2–64 or 4–64 kHz; *Dynamic moving ripple (DMR)*: a noise signal continuously modulated by an envelope defined by a smooth, random walk through septotemporal modulation space (spectral modulation range: 0–2 cycles/octave; temporal modulation range: −20–20 cycles/s; modulation depth: 40 dB; Escabí et al., 2003). The noise signal contains uniform octave band energy, created by summing multiple tone carriers (64/octave), each with randomized phase and frequencies spaced evenly on an octave scale between 2–64 or 4–64 kHz. *Random double sweep (RDS)*: two uncorrelated frequency-modulated sweeps, i.e., tones of continuously variable frequency across time (frequency range: 4–64 kHz; sweep rate range: −50–50 octaves/s; Gourévitch et al., 2015); *Clicks*: 0.1 ms positive square wave pulses; *Vocalization*: a vocalization produced by a male mouse in an isolated cage after exposure to female urine recorded with an Avisoft-Bioacoustics ultrasonic microphone (acquired at 250 kHz then down-sampled to 192 kHz); *Bandpass noise*: to avoid filter artifacts at narrow bandwidths, these sounds were created by summing many tones (2^15^) with randomized phases and frequencies equally spaced on an octave scale within a range specified by a given center frequency and bandwidth (center frequency range: 4–32 kHz; bandwidth range: 1/8–2 octaves); *Chords*: harmonic tone complexes created by summing tones (1–256 carriers/octave) with randomized phases and frequencies equally spaced on an octave scale between 2–64 kHz.

In one experiment, we presented full field visual flash (transition from black to white) on a monitor positioned ∼25 cm in front of the mouse. Auditory and visual stimuli were presented alone or together using Psychophysics Toolbox Version 3 (Kleiner et al., 2007). Monitor luminance was calibrated to 25 cd/m^2^ for 50% gray at eye position. Auditory-visual onset synchrony was calibrated within ∼1 ms using a diode to record monitor luminance values.

### Treadmill and face movement recordings

We used custom software to log treadmill movements detected by optical USB mice positioned near the surface of the treadmill as in our previous work (Bigelow et al., 2022). In most experiments, we also recorded face movements using a Mako U-130B camera (Allied Vision) and varifocal lens (B&H Photo Video) illuminated by infrared LED and positioned to include nose, mouth, whisker, and eye regions (**Figure 7A**). Frame acquisitions were triggered by an Arduino UNO at either ∼15 or ∼30 frames/s. We then used FaceMap (Syeda et al., 2024) offline to calculate face motion energy (FME) across time. This includes converting videos into difference frames, reducing dimensionality across time with PCA, and projecting individual difference frames onto principal components for a time series reflecting relative FME in arbitrary units. Outlier values were common in both treadmill and face movement recordings; we thus limited each time series between the 99 and 0.1 percentiles for the recording. We then normalized these values as percentages of the maximum within each recording.

### Analysis

In most cases, we used peristimulus-time histograms (PSTHs) to assess spiking responses to sound, with exceptions indicated below. Spiking responses in sensory areas and elsewhere have diverse temporal dynamics, featuring a wide range of latencies and durations in response to stimulus onset and/or offset, reflecting excitation and/or suppression relative to pre-stimulus firing. To accommodate this wide range of potential response patterns, we used a response reliability metric to determine whether stimulus-aligned firing was statistically significant for each unit (Bigelow et al., 2022). First, we calculated the correlation coefficient between two PSTHs constructed from random trial halves (without replacement). We then repeated the process 1000 times and defined response reliability as the mean across iterations. We further defined chance reliability in the same way after circularly shifting individual trials in time by a random value, thus breaking the temporal relationship between stimulus and response without altering other statistics such as spike counts and interspike intervals. A p-value was then calculated by z-score transforming the observed reliability value using the mean and standard deviation of the chance (shifted) distribution and identifying its associated probability in the cumulative normal distribution function. In many experiments, stimuli varied along one or more dimensions (e.g., tone frequency). In such cases, we calculated reliability separately for each stimulus value, as well as a single composite reliability value reflecting concatenated responses across all stimulus values.

We analyzed responses to continuous DMR sounds (**Figures 2 and 4**) using reverse correlation (Bigelow et al., 2022). For each unit, we calculated the spike-triggered average (STA) by summing windowed stimulus segments (time × frequency) around with each spike and dividing by spike count (**Figure 2B, i**). The STA is thus a linear estimate of the unit’s spectrotemporal receptive field (STRF). We used a temporal window spanning 200 ms before and 50 ms after each spike discretized in 5-ms bins, and a frequency window spanning the full stimulus range (2–64 kHz), discretized in 1/32 octave bands. Spikes from the first 200 ms of the stimulus were not included in STA calculations to reduce bias introduced by onset transients. We also calculated a null STA, reflecting time-frequency bins expected by chance, using the same procedure after reversing the stimulus in time. For display purposes, we transformed STA time-frequency bin values into z-scores relative to the null STA distribution, smoothed with a gaussian window (σ = 3 bins), and set values below |2.33| to zero to display only values unlikely to occur by chance (p < 0.01). We then assessed the statistical significance of the STA using the response reliability metric described above. For experiments using continuous 30-min DMR, this was implemented by discretizing the full stimulus into contiguous 1-min trials.

**Figure 2.**
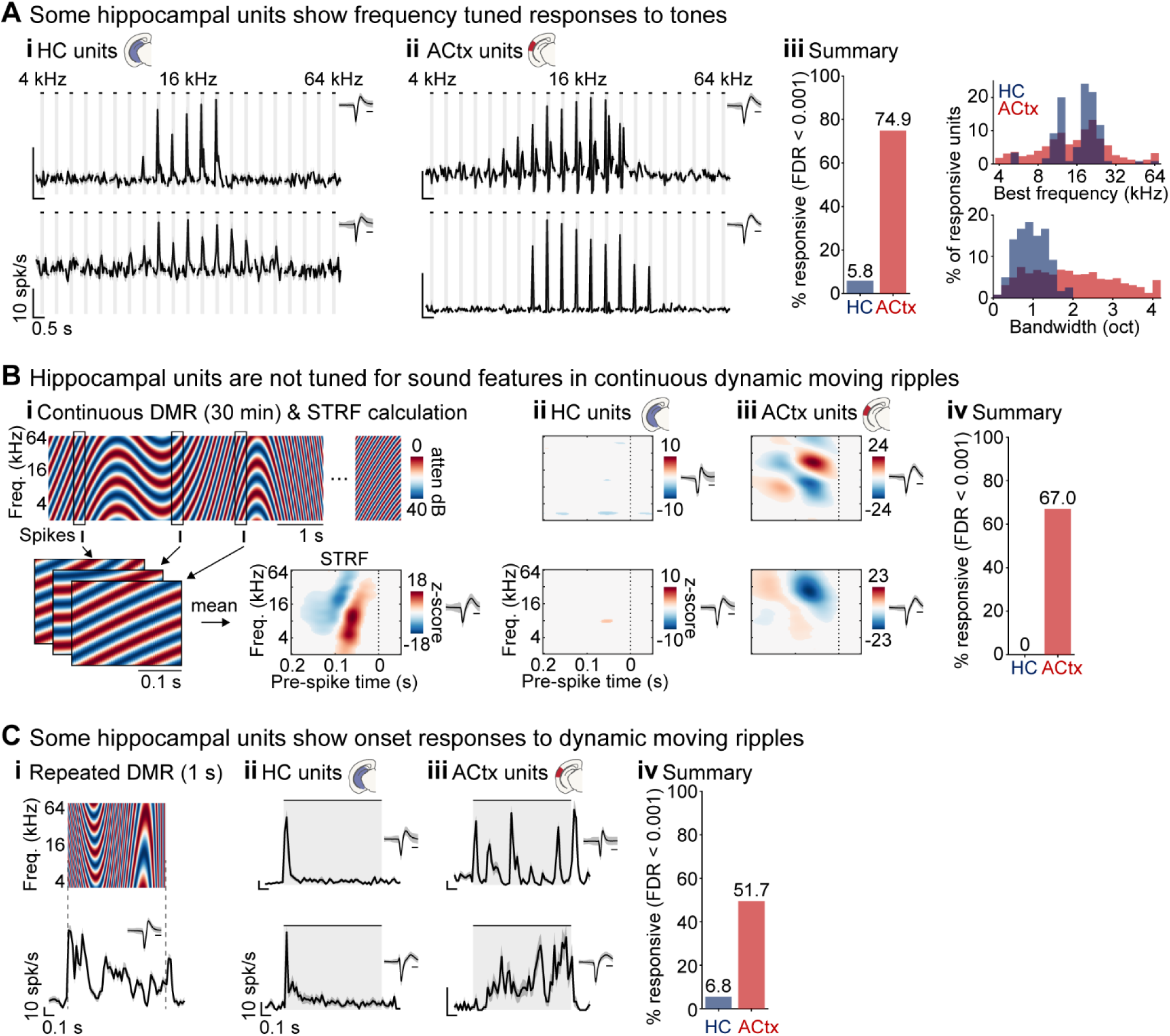
Hippocampal units show tuned responses to tone pips but not continuous dynamic moving ripples. (**A**) Some HC units show frequency tuning for tones. Example unit responses to tones in (**i**) HC and (**ii**) ACtx. (iii) Percentages of units with reliable responses to tones (left), and their corresponding best frequencies (top right) and bandwidths (bottom right). (**B**) HC units are not tuned for features in continuous DMR. (**i**) DMR stimulus and illustration of STRF calculated by averaging the stimulus segments preceding each spike for an example ACtx unit. (**ii**) Example STRFs from HCs unit lacking obvious structure (**iii**) Example ACtx STRFs. (iv) Percentages of units with reliable STRFs indicating HC units did not encode features in continuous DMR. (**C**) Some HC units responded to repeated a DMR segment. (**i**) Repeated DMR segment and example ACtx unit response. (**ii**) Example HC units showing obvious onset responses. (**iii**) Example ACtx units showing responses throughout the stimulus. (**iv**) Percentages of with reliable responses to the DMR segment.

In all analyses, we adjusted for multiple comparisons using Benjamini–Hochberg False Discovery Rate (FDR) correction applied across all recorded units for a given experiment (Benjamini & Hochberg, 1995). Except where otherwise indicated, we used a conservative α = 0.001 to identify units with highly reliable responses.

## Results

We used Neuropixels probes to simultaneously record HC and ACtx responses to a wide variety of sounds presented passively to awake, naïve mice. We cumulatively recorded from thousands of units in each region across eight separate experiments, thus enabling detailed analysis of the stimulus preferences and response characteristics of units in each area.

### Passive sounds evoke short latency responses in some hippocampal units of naïve mice

We first replicated previous findings suggesting some HC neurons show rapid responses to tones and broadband noise. Latency calculations reflect deviations from pre-stimulus activity binned at high temporal resolution and thus benefit from averaging across many trials. For tone experiments, we presented 120 repetitions each of frequencies spanning 4–64 kHz in 0.2 octave steps (150 ms duration, 70 dB SPL). We then constructed frequency-averaged PSTHs for each unit using 2-ms bins, smoothed with a Savitzky-Golay filter (3rd order, 10 ms window). Onset latency was defined as the first bin for which activity fell above or below pre-stimulus firing (mean ±3 SD) for at least three consecutive bins (**Figure 1D**; Morrill & Hasenstaub, 2018). We recorded from a total of 2,128 HC units and 2,263 ACtx units across 37 recordings from 17 mice (12 female). Of these, 124 HC units and 1,493 ACtx units had measurable response latencies within the stimulus window (offset responses are considered in **Figure 3**). Consistent with the short latencies reported by previous studies, HC units in our sample had latencies of 23 ±13.4 ms (median ±MAD) across excited and suppressed responses, compared to 21 ±16.7 ms for ACtx units. Excited responses typically occurred earlier than suppressed responses in both areas (HC: 23 vs. 34 ms; ACtx: 19 vs. 29 ms).

**Figure 3.**
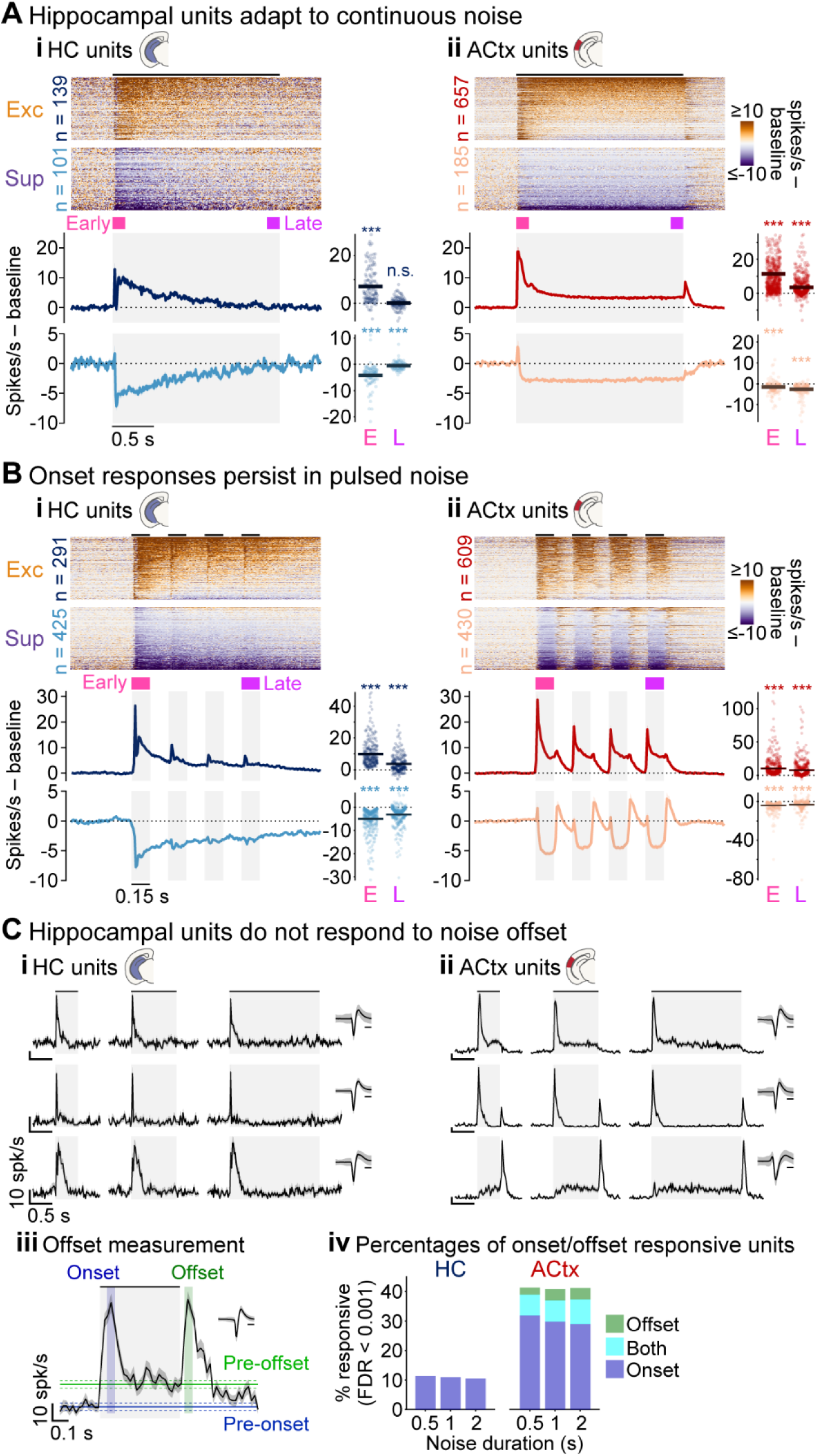
Hippocampal units respond primarily to sound onset. (**A**) HC responses adapt to 2 s broadband noise. (**i**) Individual HC unit responses (top) and means (bottom) separated into groups reflecting excitation and suppression relative to baseline firing. Population rates averaged within Early and Late windows (right) indicated activity returned near baseline levels by the end of the stimulus. (**ii**) ACtx unit responses organized as in (i). (**B**) Onset responses persist in pulse noise. (**i**) HC unit responses (top) and means (bottom) as in (Ai). Window averaged rates (right) indicated onset responses remained different from baseline through the final pulse. (**ii**) ACtx responses organized as in (i). (**C**) HC units do not respond to noise offset. (**i**) Example HC units showing responses at noise onset only. (**ii**) Example ACtx units with responses at noise onset and/or offset. (**iii**) Onsets and offsets were measured within 50-ms windows surrounding the peak and compared to an equivalent pre-stimulus window. Offset responses were also compared to a pre-offset window to ensure did not reflect sustained responses during the sound period (**iv**) Percentages of units with responses at noise onset only, offset only, or both, indicating a lack of offset responses in HC units.

We similarly calculated noise response latencies in separate recordings presenting 500 repetitions of a four-pulse broadband noise train (white noise filtered with a 4–64 kHz passband; 150 ms pulse, 150 ms inter-pulse interval; 65 dB SPL; 3 s inter-trial interval; Olsen & Hasenstaub, 2022). We measured onset latencies within the first 150 ms pulse in 938 of 2,648 HC units (35.4%) and 1,666 of 3,152 ACtx units (52.8%) recorded in 31 experiments from 16 mice (6 female). Similar to tones, median response latencies for noise were 25 ±15.2 ms in HC compared to 17 ±20.2 ms in ACtx, with most excited responses appearing first (HC: 21 vs. 29 ms; ACtx: 17 vs. 29 ms). We explored the finding that many more HC units responded to the noise train than tones in additional experiments below (**Figures 4–5**).

**Figure 4.**
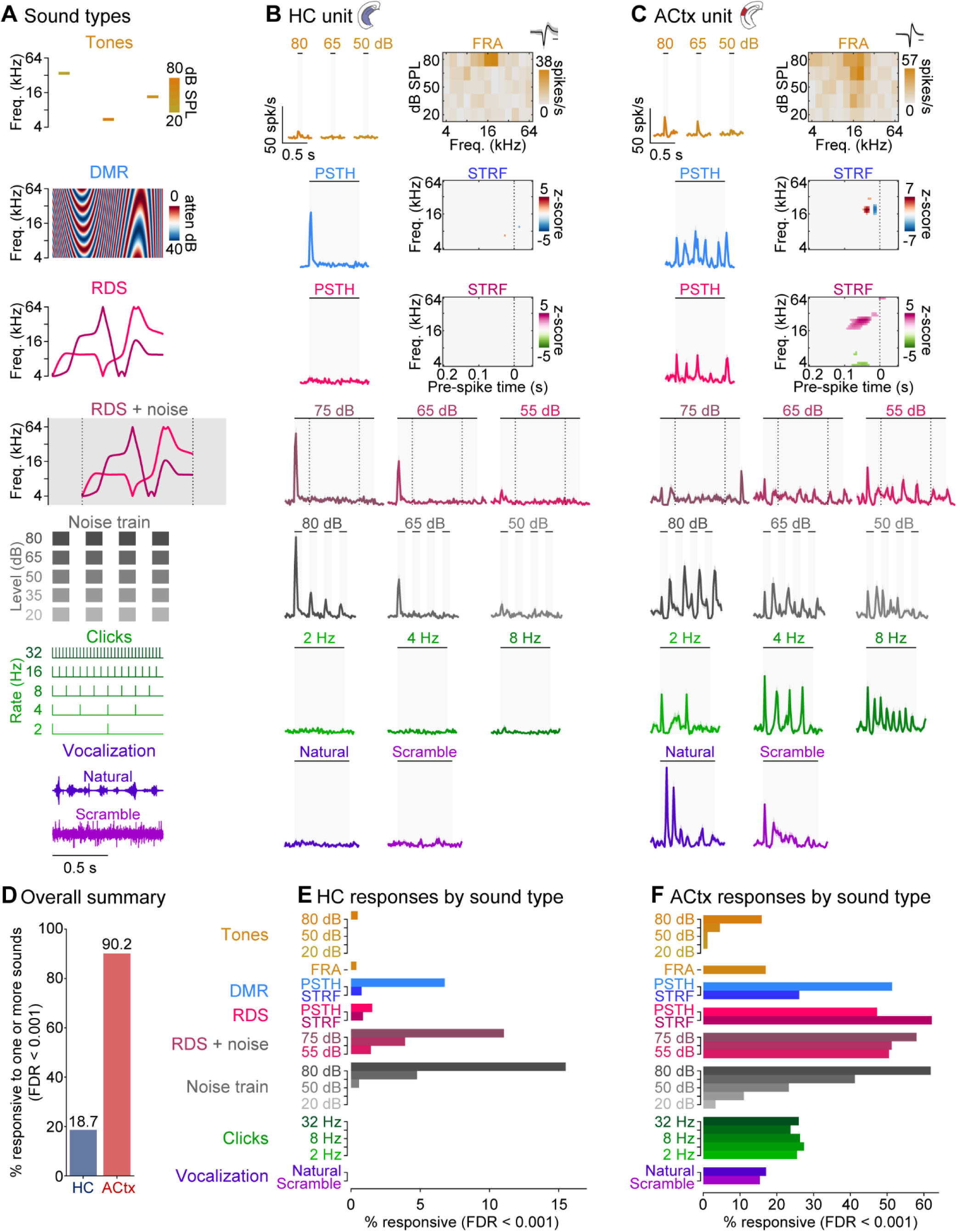
Hippocampal units respond best to broadband noise among diverse sound types. (**A**) Diverse sound types interleaved in pseudorandom order within the same experiment. (**B**) Example HC unit showing responses to RDS + noise, noise train, DMR, and 80 dB tones. (**C**) Example ACtx unit responding to all tested sound types. (**D**) Percentages of units with responses to one or more of the tested sounds. (**E**) HC units primarily responded to sounds that included broadband noise, especially the Noise train and RDS + noise stimuli. (**F**) ACtx units responded to all sound types.

**Figure 5.**
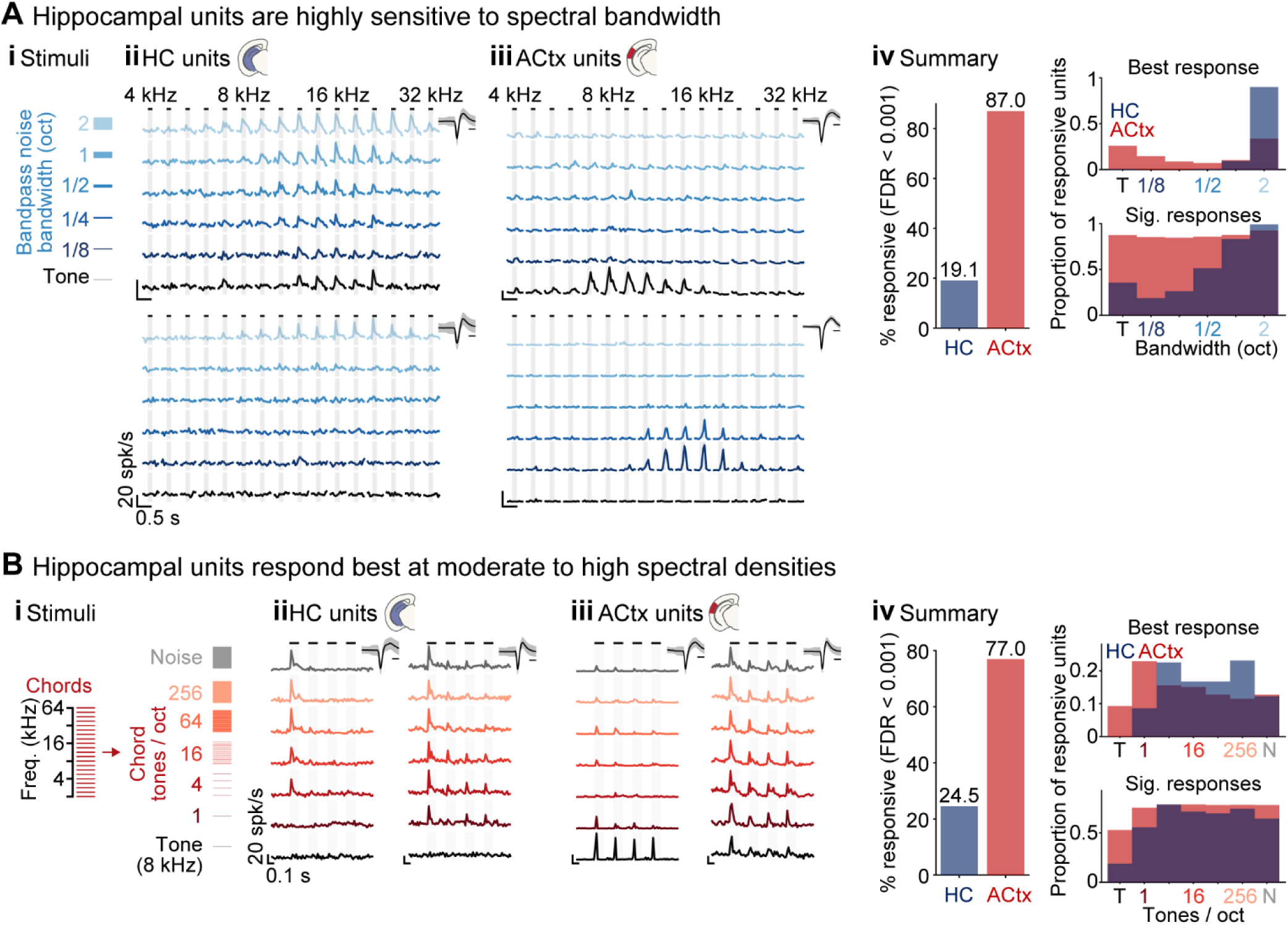
Hippocampal preference for noise reflects spectral bandwidth and density. (**A**) HC units are highly sensitive to spectral bandwidth. (**i**) Stimuli comprised tones and bandpass noise with bandwidths ranging from 1/8 to 2 octaves. (**ii**) Example HC units with strong responses to 2 octave noise. (**iii**) Example ACtx units selective for tones (top) and narrowband noise (bottom). (**iv**) Percentages of units with reliable responses (left), and the bandwidths eliciting the best response (top right) and significant responses (bottom right). (**B**) HC units are sensitive to spectral density. (**i**) Chord stimuli comprising 1 to 256 tones/octave. (**ii**) Example HC units responsive to chords but not tones. (**iii**) Example ACtx units with strong responses to tones (left) and all tested sounds (right). (**iv**) Percentages of units with reliable responses (left), and the sound types eliciting the best response (top right) and significant responses (bottom right).

### Hippocampus shows frequency tuning for tone pips but not continuous dynamic moving ripples

An obvious question regarding tone responses in HC is whether they are frequency tuned. If so, how do their tuning characteristics compare with ACtx? Might they be tuned for other sound features that drive selective firing in ACtx units? We addressed these questions by first examining tone frequencies that evoked responses in each area. Of 2,128 HC units, 124 had significant composite response reliability (5.8%) compared to 1,695 of 2,263 in ACtx (74.9%). Example units in **Figure 2A** show responses with clearly limited frequency ranges in each area. Best frequencies in HC units (the frequency with maximum response reliability) clustered almost exclusively between 8 and 32 kHz, corresponding to the most sensitive hearing range in mice (**Figure 2Aiii, top**). Best frequencies for ACtx units were similarly most common between 8 and 32 kHz but also covered the octaves above and below. Total responsive bandwidth (all tone frequencies with reliability p < 0.05) was 1 ±0.32 ms in HC (median ±MAD) compared to 2 ±0.9 ms in ACtx, a significant difference (p < 10^−33^, Wilcoxon rank sum test). Thus, some HC units are tuned for tone frequency with more stereotypical and narrow frequency preferences than ACtx units.

Would auditory tuning observed for tone frequency generalize to other sound features? Auditory neurons show selectivity for many sound features including modulations in the spectral and temporal domains (Atencio & Schreiner, 2008). A standard approach for measuring these preferences is presenting continuous DMR and averaging across the stimulus segments preceding each spike (Atencio & Schreiner, n.d.), i.e., the STA (**Figure 2Bi**). We calculated STAs for each HC and ACtx unit in a dataset of 73 recordings from 29 mice (17 female). Consistent with previous studies, the majority of ACtx units had reliable STAs (3,159 of 4,716; 67.0%). In contrast, not a single HC unit out of 3,829 had a reliable STA (**Figure 2Biv**). Thus, HC tuning was sparse, stereotypical, and narrow for tone pips but non-existent for continuous DMR.

By design, continuous DMR lacked recurring silent interstimulus intervals and sound onsets present in the tone and noise train experiments. We speculated HC responses in those experiments may have depended on transitions from silence to sound. To test this idea, we analyzed responses to 50 repetitions of a 1-s DMR separated by 300 ms of silence, which were interleaved with other sound types in a separate experiment described further below (Figure 4). Across 26 recordings in 11 mice (9 female), we found 71 of 1,051 HC units (6.8%) responded to this version of DMR (**Figure 2Civ**) – comparable to the percentage responsive to tones (Figure 2Aiii). These findings suggested sound onsets may be critical for HC responses.

### Hippocampus primarily responds to sound onset

We conducted three additional analyses aimed at clarifying whether the predominance of onset responses in HC might reflect adaptation to sounds with stationary stimulus statistics, and whether transitions from sound to silence (offsets) might be similarly effective in activating HC units as transitions from silence to sound (onsets). We first examined activity evoked by 2 s continuous broadband noise segments separated by 1.4–1.5 s silent intervals in an experiment presenting 0.5, 1, and 2 s segments 50 times each in random order (4–64 kHz bandpass filtered white noise; 65 dB SPL). This dataset included 883 HC and 1,347 ACtx units from 22 recordings in 5 mice (0 female). We further analyzed units with significant increases or decreases in mean firing during the stimulus window relative to pre-stimulus firing (p < 0.001, Wilcoxon signed rank tests), reflecting 240 units from HC (27.2%) and 842 from ACtx (62.5%). As seen in **Figure 3Ai**, HC units with both excited and suppressed responses showed obvious onset responses but typically returned to baseline firing before stimulus end. Wilcoxon signed rank tests confirmed activity in excited HC units differed significantly from baseline within the first 150 ms of the stimulus (“Early” window; p < 10^−20^) but not the last 150 ms (“Late” window; p = 0.47). Similarly, suppressed HC units had large changes from baseline in the Early window (p < 10^−20^), and only subtle but significant changes in the late window (p < 10^−4^). By contrast, responses in many ACtx units were sustained throughout the entire stimulus window, giving rise to highly significant group differences from baseline across response types and time windows (all p-values < 10^− 18^). Thus, HC units respond robustly at noise onset but adapt quickly to continued noise.

Would HC units similarly adapt to a repetitive pattern of alternating noise and silence? We reanalyzed responses to the noise train stimuli used for latency calculations above described above using the same Early and Late firing windows described in Figure 3A. Onset transients diminished following the initial pulse but remained clear in HC population-averaged firing (**Figure 3B**). This was confirmed by significant Wilcoxon signed rank tests for each response type, firing window, and area (all p-values < 10^−29^). Thus, even brief silence (150 ms) is sufficient for onset responses in HC.

Sound offset responses are common throughout the auditory pathway and believed important for processing temporal features in sound (Malone et al., 2015; Olsen & Hasenstaub, 2022). As expected, responses at noise offset were apparent in many of ACtx units in Figures 3A and 3B. We quantified such responses using continuous noise of three durations (0.5, 1, and 2 s). We analyzed only excitatory responses since offset suppression is difficult to disambiguate from post-stimulus recovery effects (Kopp-Scheinpflug et al., 2018). To accommodate potentially different onset and offset latencies, we first searched 100 ms following noise onset and offset in 1 ms steps for the 50 ms window that maximized firing. We then considered onset responses significant that differed from and equivalent baseline window immediately preceding the noise (p < 0.001; Wilcoxon signed rank tests). Offset responses were analyzed similarly, except were required to differ from two baselines, one before noise onset and the other before noise offset, to ensure elevated firing did not merely reflect continuation of sustained activity during the stimulus period (**Figure 3Ciii**, Bigelow et al., 2022; Olsen & Hasenstaub, 2022). Using these criteria, we observed offset responses in roughly one of five ACtx neurons, but no HC units (**Figure 3Civ**). Thus, the onset of a sound – but neither its continuation nor offset – is relevant to HC.

### Hippocampus responds best to broadband noise among diverse sound types

Percentages of responsive HC units ranged widely among separate experiments testing different sound types above; 5.6% responded to tones (**Figure 2A**), 5.4% to repeated DMR segments (**Figure 2C**), and 27.2% to broadband noise segments (**Figure 3A**). We also measured onset responses in 35.4% of units using a broadband noise train repeated 500 times (**Figure 1C**). These differences may be partly influenced by experimenter choices such as stimulus repetitions and interstimulus intervals but also suggest differences due to acoustic features. Diverse search stimuli are often used when probing auditory pathway stations because individual neurons often respond robustly to one sound type or feature but not others. We thus examined whether HC units might show similar preferences among a battery of diverse sound types interleaved within the same experiment.

The battery was designed to elicit responses in as many ACtx neurons as possible, and included the following sounds depicted from top to bottom in **Figure 4A**: *Tones:* frequencies 4–64 kHz (1/3 octave steps), levels 20–80 dB (15 dB steps), 100 ms duration; *DMR:* frequency range 4–64 kHz, spectral modulations 0–2 cycles/octave, temporal modulations −20–20 cycles/s, 1 s repeating segments and 6 s non-repeating segments used for PSTH and STRF analysis, respectively; *RDS:* frequency range 4–64 kHz, sweep rates −50–50 octaves/s, 1 s repeating segments and 6 s non repeating segments used for PSTH and STRF analysis, respectively; *RDS + noise:* the same 1 s repeating segment presented with 1.6 s broadband noise at 55, 65, 75 dB (white noise filtered with a passband spanning 1/2 octave above and below the 4–64 kHz RDS frequency range); *Noise train:* four-pulse broadband noise train presented at levels spanning 20–80 dB (15 dB steps; white noise filtered with a 4–64 kHz passband; 150 ms pulse, 150 ms inter-pulse interval); *Click trains:* 0.1 ms square wave pulses presented at rates 2–32 Hz (octave spacing); *Vocalization:* natural mouse vocalization presented as it was recorded (natural) and after randomizing the samples in time (scrambled). All sounds were interleaved in random order with each sound repeated 50 times (300 ms interstimulus intervals) except tone pips which were repeated 12 times per frequency-level combination. Except where otherwise indicated, all sounds were calibrated to 65 dB SPL. Responses to non-repeating DMR and RDS segments were analyzed using STA reliability. Tone responses were analyzed two ways. First, frequency response area (FRA) functions were constructed by averaging firing within 50 ms of tone onset for each frequency-level combination. Response reliability was then calculated as for other analyses by subsampling random trial halves. Second, frequency-averaged PSTHs were constructed for each level and assessed by response reliability. Responses to all other sounds were analyzed using PSTH response reliability.

As expected, the vast majority of ACtx units (1,494 of 1,657; 90.2%) had reliable responses to at least one sound in this battery (**Figure 4D**). Many HC units also responded to at least one sound (196 of 1,051; 18.7%). Parsing these responses by sound type revealed HC units primarily responded to sounds with broadband noise characteristics (**Figure 4E**); few to none responded to sounds other than noise train, RDS + noise, and repeated DMR. This pattern was not observed for ACtx units, which responded to all sounds and especially well to RDS and DMR (**Figure 4F**). Surprisingly, no HC units responded to the brief broadband signal generated by 0.1 ms click pulses, even when presented at a comparable repetition rate to the pulsed noise train (4 Hz). Click responses were robust in many ACtx units, suggesting differences between the reticular-limbic and canonical auditory pathways in the way acoustic energy is integrated across time. Because loudness percepts correlate with sound duration (Clayton, McGill, et al., 2024), this could be related to the higher sound level thresholds in reticular-limbic stations (Xiao et al., 2018).

### Hippocampal responses to broadband noise reflect spectral bandwidth and density

Broadband noise was clearly the most effective stimulus for activating HC units in the experiments above. But which acoustic features explain its effectiveness? What makes noise noisy? We contrasted responses to noise and tones by manipulating two parameters: spectral bandwidth and density.

To assess spectral bandwidth, we presented tones and bandpass noise with bandwidths ranging from 1/8 to 2 octaves (octave spacing) and center frequencies spanning 4–32 kHz (0.2 octave spacing; 150 ms duration). Sound levels were equated for all stimuli at 65 dB SPL and each frequency-bandwidth combination was repeated 50 times. Many units in HC (279 of 1,463; 19.1%) and most in ACtx (1,687 of 1,940; 87.0%) had significant composite response reliability for this stimulus set (**Figure 5Aiv**). HC responses were indeed strongly dependent on bandwidth, with nearly all units responding best to 2 octave stimuli (**Figure 5Aiv, top**; measured by composite reliability across frequency within bandwidth). This distribution was significantly different from ACtx (p < 10^−66^, Wilcoxon rank sum test), where many units preferred either tones or 2 octave noise, and some preferred intermediate bandwidths.

We assessed spectral density using chord analogs of the four-pulse noise train used for experiments above, comprising harmonic tone complexes constructed from 1 to 256 carriers/octave (**Figure 5Bi**). The chords thus had equivalent bandwidth but variable density. We also included broadband noise and tone trains (8 kHz) for comparison. We equated all sound levels at 65 dB SPL and repeated each stimulus 50 times. Similar to bandpass noise, this stimulus set evoked responses in many HC units (359 of 1,463; 24.5%) and most ACtx units (1,493 of 1,940; 77.0%), measured by composite response reliability (**Figure 5Biv**). Population analysis indicated HC units responded best with 4 or more carriers/octave (**Figure 5Biv, top**), again reflecting preference for noisier sounds relative to ACtx units (p < 10^−12^, Wilcoxon rank sum test).

### Hippocampal responses to noise are more prevalent than to visual flash

The experiments above indicated most responses to passive sound in HC reflect the sudden occurrence of noise in an otherwise quiet environment. These outcomes suggested units with such responses may be involved more generally in environmental change detection. If so, they might respond to other environmental changes including visual events. We tested this possibility with visual flash trains similarly structured to the noise train stimuli described above (150 ms flash, 150 ms inter-flash interval). Noise and flash trains were presented alone or together with 50–200 repetitions of each condition randomly interleaved with 2–3 s intertrial intervals. Roughly half of HC units in this sample responded to the noise train (**Figure 6C, left**; 868 of 1,604; 54.1%). Responses to flash alone were far less common (202 of 1,604; 12.6%), suggesting HC units may be especially sensitive to changes in the acoustic environment. Most flash-responsive units also responded to noise (**Figure 6C, right**), suggesting audiovisual convergence in some HC units. Consistent with this finding, more units responded to noise-flash trains presented together than either alone (1,022 of 1,604; 63.7%), suggesting multisensory facilitation. The percentage of flash responsive units was similar in ACtx (350 of 2,274; 15.4%), and nearly all ACtx units responded to noise (2,140 of 2,274; 94.1%).

**Figure 6.**
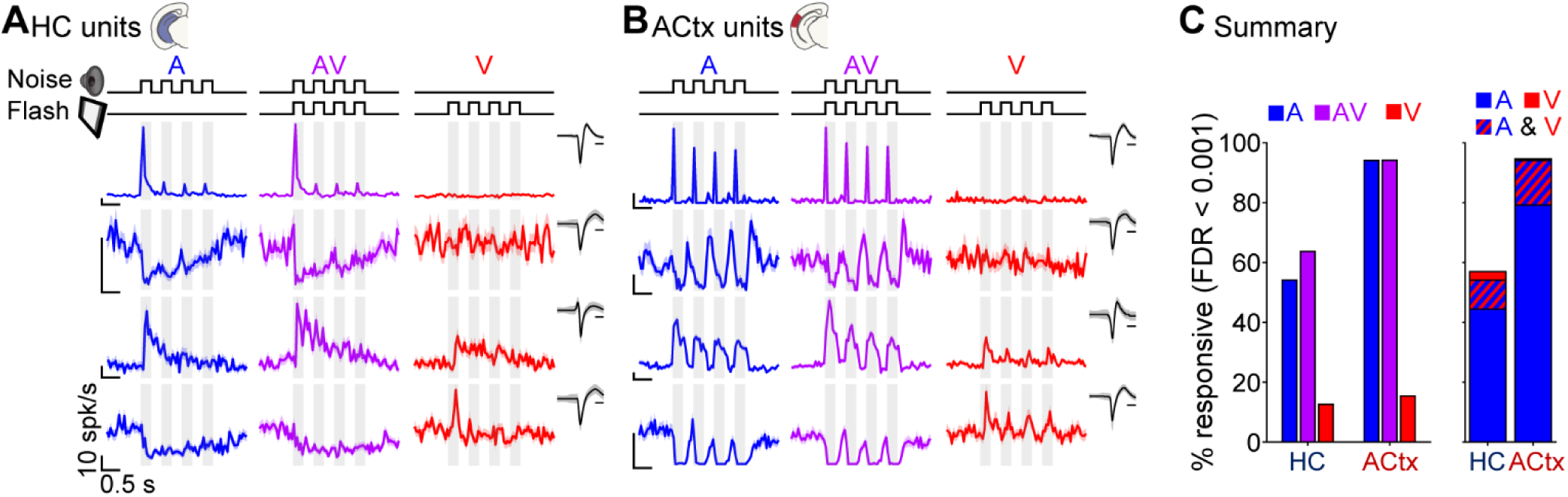
Hippocampal responses are more prevalent for noise than visual flash. (**A**) Example HC units responsive to noise only (top two units) or both noise and flash (bottom units). (**B**) Example ACtx units as in (A). (**C**) Percentages of units with reliable responses to noise and/or flash trains indicating that responses to noise are more prevalent in HC units (left) and most flash responsive units also respond to noise (right).

### Hippocampal responses to sound correlate with face movement

Recent studies from independent groups have reported that sounds can evoke face movement responses, which may correlate with certain aspects of the sound. Two findings are especially relevant to our study. First, face movement evoked by natural sounds correlated strongly with population activity in HC (Bimbard et al., 2023). Second, face movement responses were stronger for broadband noise than tones or visual stimuli (Clayton, Stecyk, et al., 2024; Olsen & Hasenstaub, 2025). We thus tested whether the prevalence of HC responses to noise vs. tones or visual flash in our study was related to stimulus evoked face movement.

We used FaceMap (Syeda et al., 2024) to transform our face video recordings into FME signals (**Figure 7A**). Consistent with previous studies, we observed clear increases in FME aligned to sound onsets in some trials (e.g., example trace with repeated noise train in **Figure 7Aii**). Trials with evoked face movements were observed to be interspersed among others without obvious FME responses to the same sounds. We also observed transient FME increases during some silent intertrial intervals. These findings replicated previous outcomes suggesting sounds can elicit FME responses with a high degree of trial variability (Olsen & Hasenstaub, 2025).

**Figure 7.**
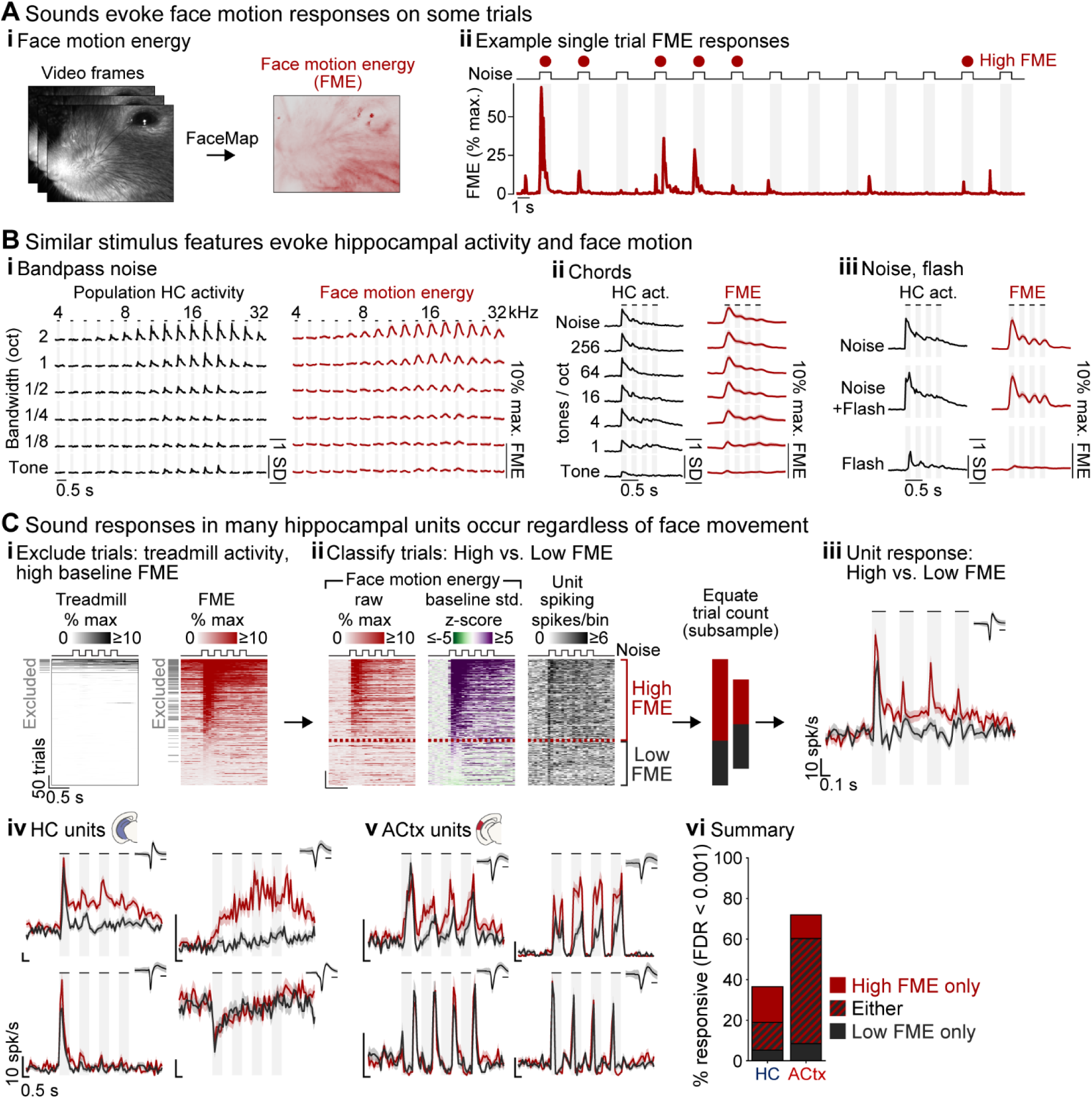
Hippocampal responses to sound correlate with face movement. (**A**) Sounds evoke face motion responses on some trials. (**i**) Raw video frames are converted into face motion energy (FME) signals using FaceMap. (**ii**) Example single trial FME responses to a broadband noise train illustrating high trial-to-trial variability. (**B**) Similar stimulus features evoke HC activity and FME. Similar patterns of HC activity and FME responses were evoked by (**i**) bandpass noise, (**ii**) chords, and (**iii**) noise/flash train stimuli. (**C**) Sound responses in many HC units occur regardless of face movement. (**i**) Trials with high baseline FME and treadmill movement were excluded to isolate stimulus-evoked FME. (**ii**) Trials were classified as High or Low FME (cutoff: 2 SD change from baseline) then equated via subsampling prior to (**iii**) averaging spiking responses. Example units showing diverse dependence on FME in both (**iv**) HC units showing diverse and (v) ACtx. (**vi**) Percentages of units with reliable sound responses on trials with High FME only, Low FME only, or both.

To determine whether FME responses were related to acoustic parameters predicting HC activation, we calculated average peri-stimulus FME for each of the stimuli presented in the bandpass noise, chord, and visual flash experiments above. We then defined a global population activity index in HC reflecting the absolute value of z-scored activity relative to pre-stimulus baseline, averaged across all responsive units. As seen in **Figure 7B**, FME responses were very tightly correlated with population HC activity across experiments and stimulus features.

Considering identical sounds evoked highly variable FME responses across trials (**Figure 7Aii**), we wondered whether responses of individual HC units might be predicted by these outcomes. We therefore reanalyzed responses in the experiment described above comprising 500 noise train repetitions. We were primarily interested in the relationship between stimulus-evoked FME and HC activity and thus excluded trials with high spontaneous FME during intertrial intervals. Treadmill recordings obtained in most experiments (27/31) indicated movements were typically associated with, but not necessary for elevated FME. We therefore excluded any trial for which mean treadmill activity or FME within a 1 s window prior to stimulus onset exceeded 5% of the maximum for the recording (**Figure 7Ci**). We then transformed FME values for each trial into z-scores relative to baseline (500 ms pre-stimulus window; **Figure 7Ci**). Trials for which mean z-scored FME was greater than two within the first 500 ms of stimulus onset were classified as high FME (Olsen & Hasenstaub, 2025). We then constructed PSTHs from high and low FME trials and calculated response reliability as in previous analyses (**Figure 7Ciii**). Because reliability and other spiking measures are affected by trial counts, we first equated trial counts by random subsampling from the condition with more trials. Experiments with at least 20 trials per condition were included in the final analysis, reducing the original sample to 2,041 HC units and 2,293 ACtx units from 22 recordings in 11 mice (5 female).

Example units in **Figure 7Civ** illustrate that FME correlated responses were highly variable among HC units. In some cases, firing was substantially elevated on high FME trials, especially following the initial pulse in the noise train (top row). Responses in other units showed no obvious relation to FME (bottom row). This variability is reprised by group data **Figure 7Cvi**, indicating many units responded reliably on high FME trials only (17.6%), fewer units on low FME trials only (5.2%), and many either way (13.7%). Similar observations held for ACtx units, although a larger proportion responded regardless of FME (>50%). Together, these results confirm previous findings that population HC responses correlate with FME, reveal substantial variability among individual neurons and trials, and clarify that many individual units respond to sound regardless of FME.

## Discussion

Previous studies established some HC units respond to sound outside learning and behavioral contexts. However, these studies left open questions about specific sound features capable of driving HC responses, and thus, the prevalence of such responses and how they might compare to responses in auditory pathway stations. Here we made significant progress toward answering these questions by testing a wide range of sound properties while recording from large unit samples in HC and ACtx. We found a significant minority of HC units responded to tone pips (∼5%; **Figure 2**). Many more responded to broadband noise (∼30–50%; **Figures 1, 3, 4, 6**), and intermediate percentages responded to bandpass noise and chords, which have acoustic properties intermediate between tones and broadband noise (∼20–25%; **Figure 5**). These responses invariably occurred at stimulus onset (**Figure 3**), trailed responses in ACtx by just a few milliseconds (**Figure 1D**), and reflected either excitation or suppression (**Figures 1, 3**). Unlike ACtx, HC units did not reliably encode spectrotemporal features in continuous DMR or RDS (**Figures 2, 4**). They did not respond continuously to ongoing broadband noise, or to noise offset (**Figure 3**). Despite their preference for broadband noise onsets, HC units did not respond to the broadband but brief signal generated by clicks (**Figure 4**). In summary, the most effective acoustic event for evoking HC responses in naïve mice was a transition from silence to broadband noise.

The prevalence of responses to passive environmental sounds suggests substantial innervation by auditory inputs in HC. The clear preference of HC units for broadband noise is consistent with previously reported selectivity for noise over tones in reticular-limbic pathway stations including pontine nuclei, medial septum, and entorhinal cortex (Xiao et al., 2018; G.-W. Zhang et al., 2018). Why do broadband noise events differentially activate limbic circuitry? One possibility is that noisy sounds may signal potential danger, such as movement by a predator or conspecific rival. In many species, vocalizations signaling aggression have noisier acoustics than affiliative calls, which are typically defined by harmonics or frequency modulation (e.g., hiss vs. meow). From this perspective, the traditional view that auditory neurons in HC are primarily involved in behavioral responses is not necessarily in conflict with the finding that they also respond to passive environmental sounds; the short latency onset transients evoked by these sounds could reflect a limbic signaling cascade initiating a behavioral response to abrupt environmental change. The auditory modality may be especially important for this process, considering responses were severalfold more common for sounds than analogous visual events (**Figure 6**).

Are HC responses to sound mediated by auditory inputs? Our results confirmed previous findings that some sounds – especially broadband noise – evoke face movement responses that correlate with population activity in HC. Thus, sound-aligned HC activity could reflect auditory and/or motor inputs. We found units consistent with each of these possibilities. We first observed that sound evoked FME is highly variable; movement responses to identical sounds ranged from strong to indiscernible on trials separated by only a few seconds (**Figure 7Aii**). Roughly 40% of apparently sound-responsive units responded only on trials with high FME, as would be expected of units that were driven by sound-evoked facial movements but not by sounds themselves. But roughly 60% of units showed the opposite pattern, responding to sounds independent of face movement or only on trials without face movement, as would be expected of units that were genuinely sound-driven (**Figure 7Cvi**). Strong face movements were thus not required for sound responses in many or most HC units. This finding is consistent with the short auditory response latencies of most HC units (**Figure 1**), as well as the candidate reticular-limbic sources for these responses (Xiao et al., 2018; G.-W. Zhang et al., 2018). Collectively, our results suggest sound responses in HC may reflect a complex volley of inputs including a rapid auditory-mediated component. Future work clarifying the circuitry underlying sound evoked face movement could be helpful for disambiguating the balance and temporal sequence of auditory-motor inputs into HC as well as ACtx.

Regardless of their precise input sources and functional roles, the finding that sound responsive neurons are common in HC could be relevant to many aspects of hearing health and disease. Occupational and environmental noise hazards have well known adverse effects on auditory pathway structures, which are believed to contribute to hearing deficits such as presbycusis and hidden hearing loss (Gourévitch et al., 2014). This raises the question of whether HC neurons might similarly be affected by hazardous noise, and whether the resulting changes to their functional properties could affect general cognitive function. Consistent with this possibility, one study found rats reared in pulsed broadband noise later exhibited spatial learning and memory deficits and diminished long-term potentiation in HC (Y. Zhang et al., 2021). Thus, the adverse effects of environmental noise could extend beyond those indicated by traditional audiologic assessment.

Hearing loss has recently been identified as a risk factor for dementia including Alzheimer’s disease (AD). Because HL is both common and strongly predicts dementia, it ranks among leading risk factors in terms of total population attributable fraction (Livingston et al., 2017, 2024, 2024). The precise mechanisms mediating this risk are not well understood, and likely include complex interactions among psychosocial factors, genetics, and other variables (Griffiths et al., 2020; Nadhimi & Llano, 2021). Nevertheless, experiments in rodent models suggest hearing loss induction can induce structural and functional changes in HC (Liu et al., 2016, 2018; Qian & Ricci, 2020), including exacerbation of pathology relevant to AD (Paciello et al., 2021). Considering AD is believed to initiate in the hippocampal complex, these studies provide proof-of-concept that auditory deafferentation could directly alter hippocampal function, independent from indirect effects of reduced communication abilities or social withdrawal. Our findings add further support to this notion by revealing that HC responses to passive auditory inputs are common – much more so than passive visual inputs. Deafferenting these inputs could thus have an outsized impact on HC function. Considering AD and hearing loss are each leading causes of disability and economic burden worldwide (Nadhimi & Llano, 2021), there is need for continued research into the relationship between hearing and AD, including whether hearing assessments and interventions could improve early detection and prevention efforts.

## Acknowledgements

This work was supported by The National Institutes of Health (R01AG078132, R01DC021595, and R01NS116598), Hearing Research Inc., The Klingenstein Foundation, PBBR Breakthrough Fund, and the Coleman Memorial Fund. We thank Christoph Schreiner and Timothy Olsen for helpful comments on the manuscript, and Nerissa Hoglen for providing mouse vocalization recordings.

